# An Oscillatory Mechanism for Multi-level Storage in Short-term Memory

**DOI:** 10.1101/2021.07.29.454329

**Authors:** Kathleen P. Champion, Olivia Gozel, Benjamin S. Lankow, G. Bard Ermentrout, Mark S. Goldman

**Author notes:** **Correspondence:** Mark Goldman, Center for Neuroscience, UC Davis, 1544 Newton Court Davis, CA 95616, Bard Ermentrout, Department of Mathematics, University of Pittsburgh, Pittsburgh, PA, 15213. These authors contributed equally to this work (Kathleen P. Champion, Olivia Gozel, Benjamin S. Lankow). This work was completed outside current employment at Amazon.com, Inc (Kathleen P. Champion).

## Abstract

Oscillatory activity is commonly observed during the maintenance of information in short-term memory, but its role remains unclear. Non-oscillatory models of short-term memory storage are able to encode stimulus identity through their spatial patterns of activity, but are typically limited to either an all-or-none representation of stimulus amplitude or exhibit a biologically implausible exact-tuning condition. Here, we demonstrate a simple phase-locking mechanism by which oscillatory input enables a circuit to generate persistent or sequential activity patterns that encode information not only in their location but also in their discretely graded amplitudes.

**Significance:** A core observation in many memory systems and tasks is the presence of oscillations during memory maintenance. Here, we demonstrate a mechanism for the accumulation and storage of information in short-term memory in which oscillatory activity enables a solution to long-standing challenges in modeling the persistent neural activity underlying working memory. These challenges include the ability to encode information with low firing rates, multi-level storage of stimulus amplitude without extreme fine tuning, and multi-level storage of information in sequential activity. Altogether, this work proposes a new class of models for the storage of information in working memory, a new potential role for brain oscillations, and a novel dynamical mechanism for multi-stability.

---

The maintenance of information in short-term memory is a key component of a wide array of cognitive (1, 2) and non-cognitive (3, 4) functions. However, the biophysical mechanisms that enable memory storage over the seconds-long time scale remain unclear. Single-unit studies have demonstrated a neural correlate of memory maintenance in the persistent activation of neurons whose population activity spans the memory period (reviewed in (2, 5, 6)). Theoretical studies have shown how such persistent activity can be generated by recurrent network feedback (7–9), but simple instantiations of this idea are either implausibly sensitive to mistuning or can only maintain a single elevated firing rate that is unrealistically high (the ‘low firing rate problem’, reviewed in (4, 10)), limiting storage about a given item to a single bit (‘on’ or ‘off’) of information.

Separately, previous studies have identified distinct bands of oscillatory activity in field potential recordings and EEG during the maintenance of working memory (reviewed in (11)). Such activity can be generated through cell-intrinsic mechanisms, local circuitry, or long-range interactions (12–14). However, it remains an open question whether oscillatory activity is necessary, sufficient, or even beneficial for working memory storage. Previous work has proposed how oscillations can contribute to a variety of memory functions such as the generation or maintenance of persistent activity (15, 16); the structuring of spatial codes through frequency coupling (17); and the coordination, control, and gating of memory-related activity (18–29). By contrast, other studies have suggested that oscillations could be an epiphenomenon of other computational or network mechanisms (30–32). Here, we demonstrate a potential mechanistic role for oscillations, regardless of source or frequency, by showing how the addition of oscillatory inputs to simple recurrent feedback circuits can enable both low firing rate persistent activity and a discretely graded set of persistent firing rates that increases the information capacity of a memory network.

To illustrate the core challenges that arise when generating biologically plausible models of persistent activity, consider an idealized circuit consisting of a memory neuron (or lumped population) connected to itself through positive feedback (Fig. 1A); this basic motif of recurrent excitation is the key component of most circuit models of persistent neural activity (reviewed in (4)). This simple circuit receives a brief stimulus (Figs. 1A,C,E external input) and needs to store it through persistent activity. Stable persistent activity 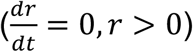 is achieved only when the intrinsic decay of the neuron (represented by the term −*r*) and the recurrent drive to the neuron (*f*(*r*)) are equal in magnitude and cancel each other. This condition imposes two separate, but related, problems that depend on whether the rate function *f*(*r*) is linear or nonlinear. In the typical nonlinear case (Fig. 1B,C), if the stimulus is too weak, the memory neuron’s low initial firing rate provides insufficient recurrent feedback to overcome the post-stimulus intrinsic decay of activity (Fig. 1B, left of open circle). As a result, the firing rate of the network returns to a low (or zero) baseline firing rate (Fig. 1C, orange, purple, green traces). By contrast, if the stimulus is stronger, the memory neuron’s initial firing provides recurrent feedback that exceeds the rate of intrinsic decay (Fig. 1B, right of open circle), leading to a reverberatory amplification of activity in which the rate rises until some saturation process brings the rate to rest at an elevated persistent level (Fig. 1C, blue and red traces). Thus, the only possibilities are that activity decays to its baseline level or that activity runs away to saturation at a high level of activity that, for typical neuronal nonlinearities, is unrealistically large. A different problem emerges in the case where the rate function *f*(*r*) is linear (Figs. 1D,E). The linearity of the rate function in this case allows a continuum of persistent rates, corresponding to the continuous set of points at which the feedback and decay lines overlap, to be stored (Fig. 1D, blue line), unlike the nonlinear case. This comes at the cost of a ‘fine-tuning’ condition: the strength of the recurrent synapse(s) must be exactly tuned to counterbalance the strength of the rate decay; an arbitrarily small violation of this condition causes the rate to exhibit runaway growth (Fig. 1E red trace) or decay to a low baseline (Fig. 1E orange trace). Although presented here for a very simple example, these problems are also commonly observed in larger neural networks (33).

**Figure 1.**
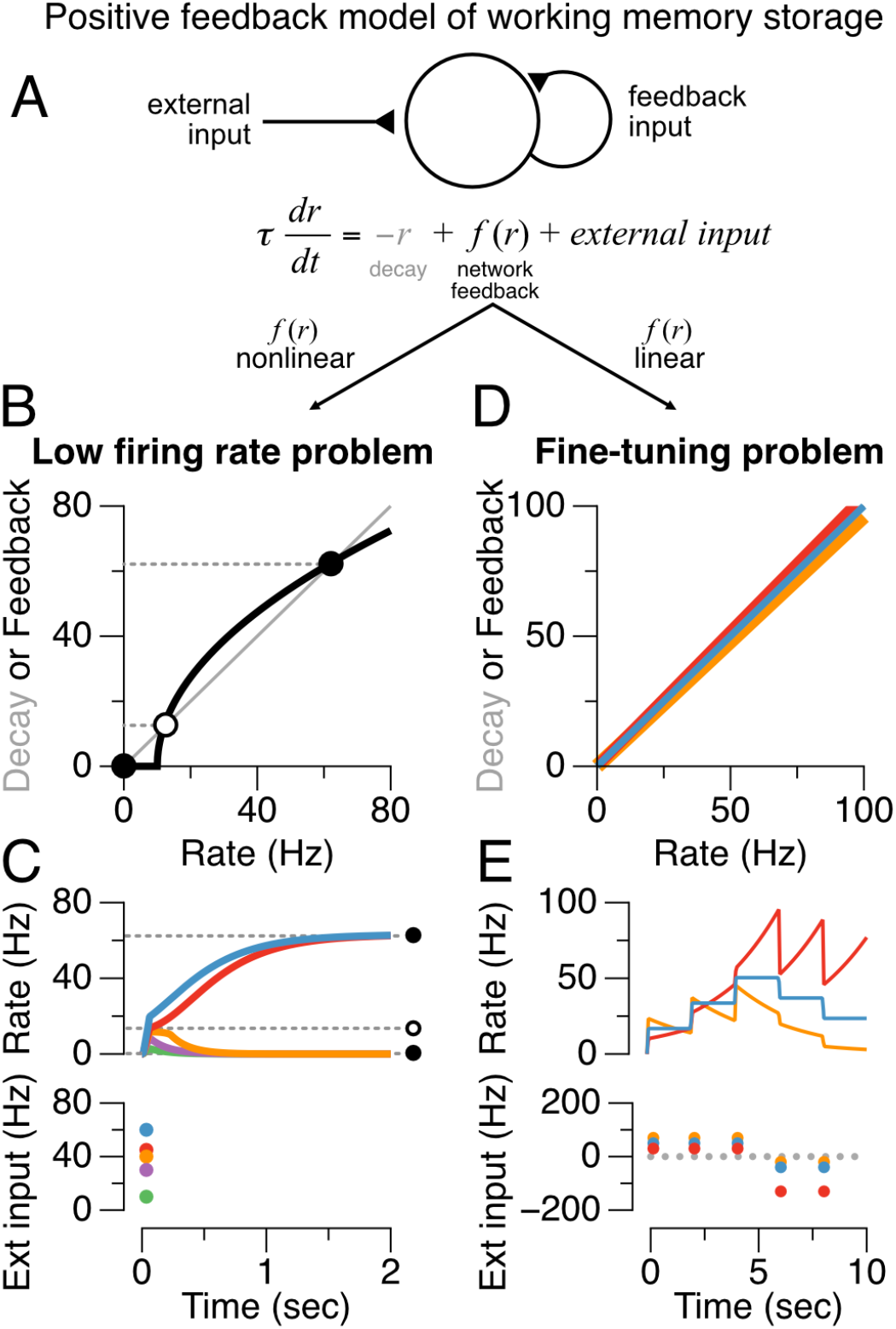
Failures of traditional positive feedback models of working memory storage. (A) Simplified model illustrating key features of positive feedback models. In the absence of external input (*external input* = 0), changes in the firing activity *r*(*t*) of a population are determined by the relative balance of network feedback (black, *f*(*r*)) and neuronal decay processes (gray, −*r*). (B,C) Nonlinear models typically exhibit a ‘low firing rate problem’. (B) During the memory period when external input is absent, the intersections of the decay (gray) and network feedback (black) functions are such that there are no stable fixed points (solid circles) within the range of firing rates typically observed during persistent neural activity. (C) Firing rates below the unstable fixed point (B,C, open circle) decay to zero (green, purple, orange lines), while firing rates above the unstable fixed point run off to uncharacteristically high rates (red, blue lines). (D,E) Linear models exhibit the ‘fine tuning problem’: minute changes in the strength of feedback (red: +5%, orange: -5%) relative to the tuned value (blue) result in instability and the inability to maintain stable persistent activity.

We next illustrate what happens when a network with the same positive feedback architecture is provided with a subthreshold oscillatory drive (Fig. 2). We demonstrate this in the more biologically realistic case of a conductance-based spiking neuron model (34) that facilitates the phase-locking phenomenon that we will describe. Without an oscillatory input, the model exhibits the ‘low firing-rate problem’ (Figs. 2A,B) and can only maintain persistent activity at a high spiking rate or not spike at all. When a subthreshold oscillation is added to the model (Figs. 2C-F), the oscillatory drive has two effects. First, it provides extra input that allows small initial inputs to trigger low-rate spiking. Second, spiking of the memory neuron does not lead to runaway feedback because, before the feedback can run away, the oscillatory drive returns towards its trough, causing a cessation of spiking. The net result is that the spike-driven feedback becomes discretized, forming a staircase whose step heights correspond to the number of spikes emitted by the neuron per oscillation cycle (Figs. 2D-F). The phase locking of the spiking to the subthreshold oscillatory drive constrains these spike numbers to be integer multiples of the oscillation.

**Figure 2.**
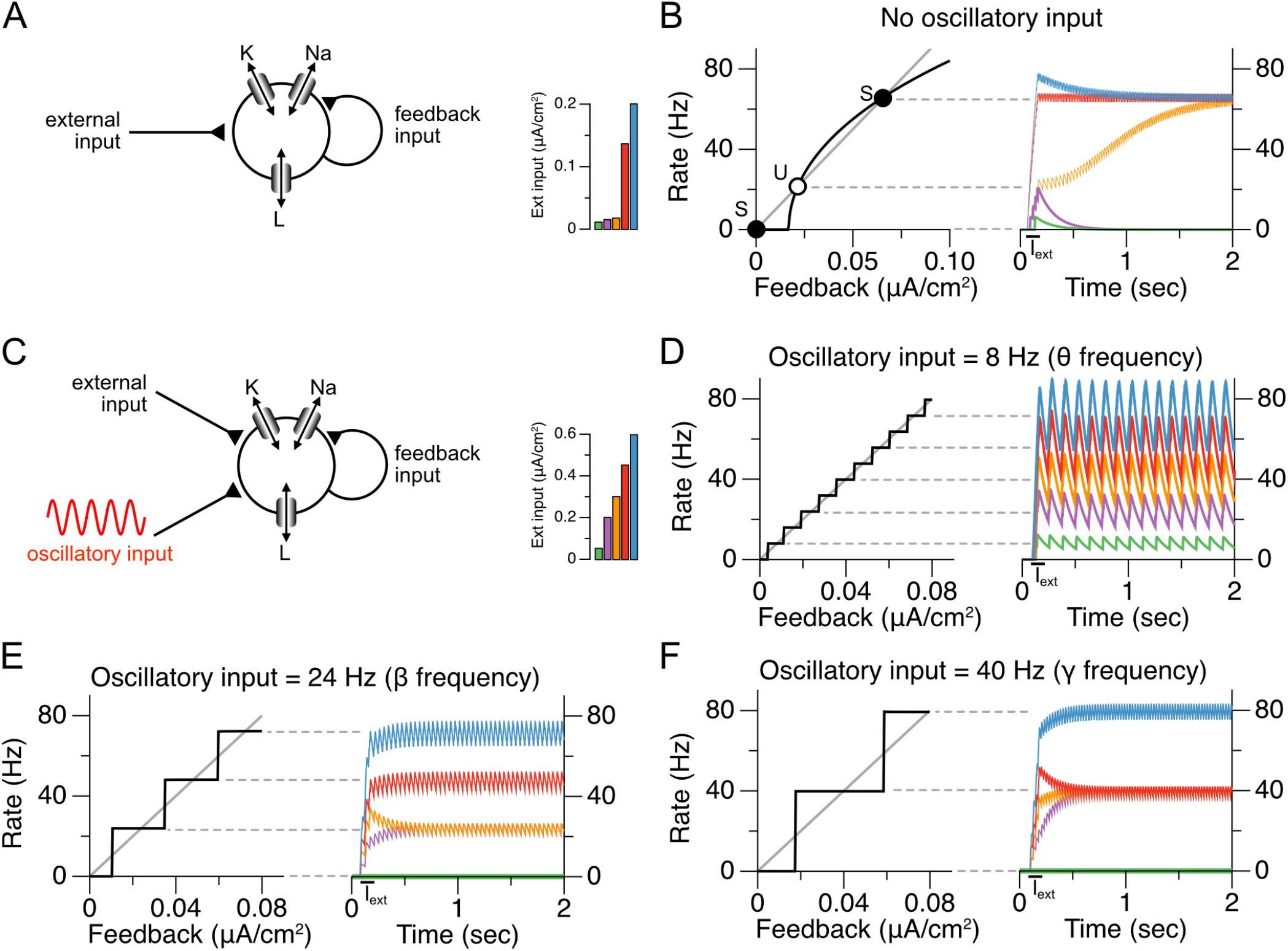
Baseline oscillatory input allows robust maintenance of discretely graded persistent activity levels in a conductance-based, spiking neuron model. (A) Schematic of conductance-based autapse model. The model is composed of potassium, sodium, and leak conductances, and receives feedback input (*I*_*syn*_ in Equation 1, Methods) as well as input current from an external source. (B) Manifestation of the ‘low-firing-rate’ problem in the conductance-based model without oscillatory input. Similar to the nonlinear firing rate model depicted in Figure 1B, the conductance-based spiking model exhibits stable fixed points only at zero and high firing rates (filled circles). Spiking rates between these two fixed points decay to zero from below the unstable fixed point (open circle) or run off to high rates from above the unstable fixed point. (C) Schematic of conductance-based neuron with the addition of an oscillatory baseline input. (D-F) Maintenance of discretely graded persistent activity levels enabled by baseline oscillatory input. Phase-locking to the oscillatory input creates stable fixed points at integer multiples of the baseline frequency. There is a trade-off between the number of firing rates that can be maintained and the robustness of these fixed points, which is related to the spacing between the fixed points. (D) Lower frequency oscillations enable a larger number of closely spaced fixed points. (E,F) Higher frequency oscillations lead to fewer, more robust, fixed points.

The key requirements for this mechanism to enable discretely graded persistent activity are the following: First, the oscillation must be strong enough to reset the activity at its troughs. Second, there must be some process that enables the activity from one cycle of the oscillation to carry through to the start of the next cycle and consequently enable renewed spiking as the oscillatory input heads towards its peak. For the simple case illustrated here, where all neurons receive oscillatory inputs that are perfectly aligned in phase, the mechanism enabling inter-cycle memory is a slow NMDA-like (or local dendritic) synaptic time constant (3, 10, 35). Alternatively, we show in the Supporting Information that, if there is heterogeneity in the phases of the oscillations received by the individual neurons in the network, the time between cycles may be bridged by the firing of other neurons in the network (Fig. S1). Some degree of tuning of the feedback is required to have multiple levels of response – such a tuning requirement is generic of models of analog or finely discretized persistent activity. In the present case, the width of the steps of the staircase provides a moderate level of robustness to mistuning, especially for higher oscillation frequencies (Figs. 2D-F). Mechanistically, this robustness occurs because errors in the tuning of feedback that are insufficient to systematically add or subtract an extra spike per cycle do not persist from cycle to cycle, unlike in models that have no oscillatory trough to reset (error-correct) the spiking activity. We illustrate this robustness to weight changes in Figure 3, where we compare the oscillatory autapse memory model (Fig. 3A,B) to an approximately linear autapse model (36, 37) that can produce (nearly) graded persistent activity (Fig. 3C,D). Each model receives an arbitrary sequence of positive and negative input pulses, and must temporally accumulate and store the pulses in persistent activity. The linear spiking autapse model requires fine tuning to maintain persistent activity: very small deviations from the tuned autapse weight lead to activity that grows to a saturating level or decays to zero activity (Figs. 3C,D). In contrast, the same synaptic weight deviations have negligible effect on the accumulation and multi-level storage capability of the nonlinear spiking neuron with oscillatory drive (Figs. 3A,B).

**Figure 3.**
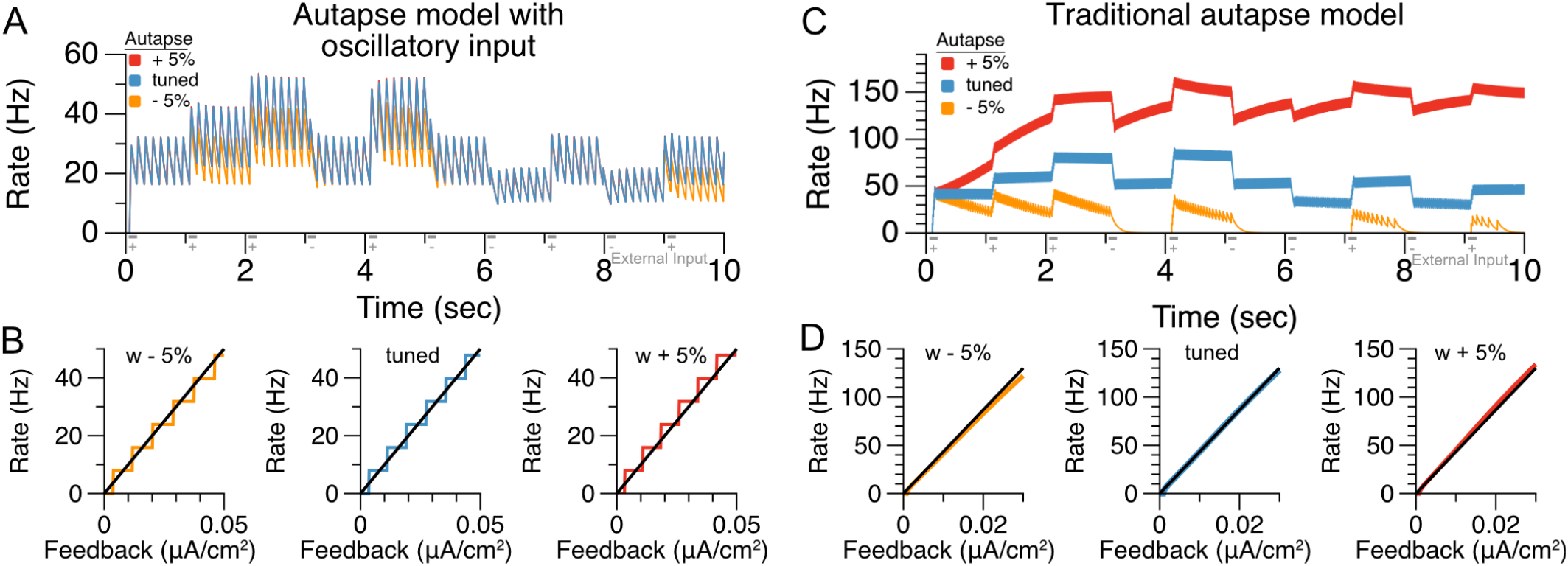
Oscillation-based integrator model exhibits more robustness to changes in the recurrent feedback weight than a traditional non-oscillation-based model. (A) Responses of oscillation-based model to a sequence of positive and negative input pulses. Red and yellow traces show the conditions in which the recurrent feedback strength has been detuned by +/- 5%, respectively. The activity levels remain persistent following detuning. (B) Steady-state firing rates as a function of synaptic activity (*I*_*syn*_ in Equation 1, Methods) that is held at steady values; mistuning the autapse strength by +/- 5% has no effect on the existence and location of the stable fixed points (intersections of black lines and horizontal stairs). (C) Responses of a traditional, approximately linear, conductance-based model of persistent neural activity (adapted from model of (37)). Detuning the recurrent feedback strength by 5% (orange and red traces) causes spiking activity to decay to 0 (orange, decreasing feedback strength) or run off to high rates (red, increasing feedback strength). (D) Small weight changes cause systematic loss of fixed points in the traditional model.

The above examples demonstrate the basic mechanism by which oscillatory input may permit discretely graded levels of firing rate to be robustly stored in a recurrent excitatory network model of persistent activity. We next explored applications of this basic principle in the case of three different network architectures: a spatially uniform (all-to-all) network that temporally integrates its inputs (Fig. 4); a ‘ring-like’ architecture whose activity can store both a spatial location and discretely graded levels at that location (Fig. 5); and a chain-like architecture that can generate sequences of activity with multiple discretely graded amplitudes (Fig. 6).

**Figure 4.**
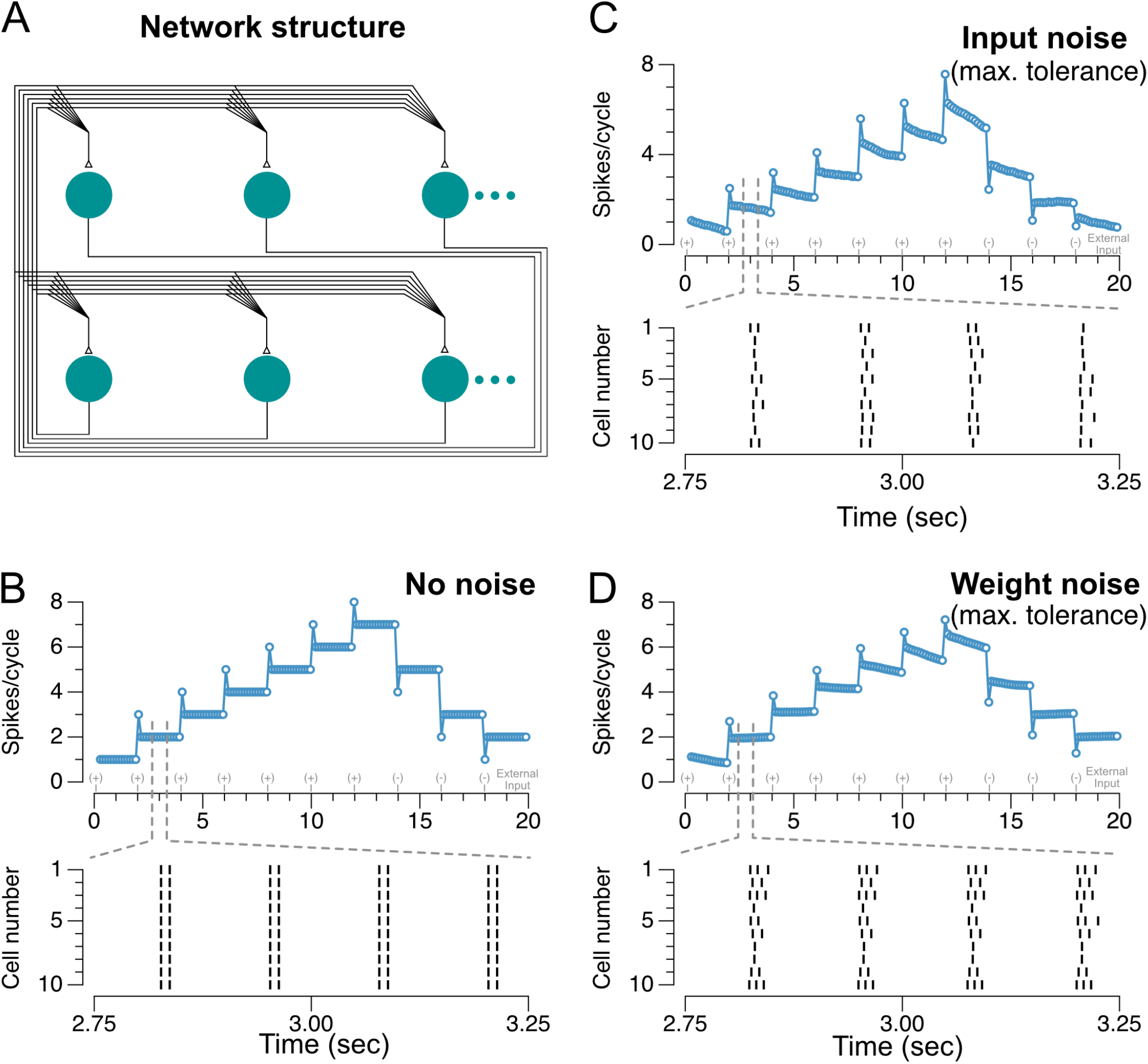
Maintenance of persistent activity and robustness to noise in a fully connected network of 1000 neurons. (A) Schematic of network. All units make synapses on all other units, with uniform synaptic weights (plus noise when present). (B) Spiking responses of network neurons to a sequence of positive and negative input pulses, with spike rasters plotted for the time window indicated by the dashed grey lines (random sample of 10 neurons from the 1000-neuron network). (C) Spiking responses of the network to the same input sequence in the presence of continuous external input noise. The noise had zero mean and standard deviation roughly one third the magnitude of the individual input pulses (*σ* = 0.05 *μA ms*^0.5^*/cm*^2^), the point at which network activity noticeably began degrading. The network is able to maintain persistent activity despite the noisy input. (D) Spiking responses of the network initialized with random noise in the connection weights. Noise with mean zero and standard deviation of 10 times the mean connectivity strength (*σ* = 0.055 *μA/cm*^2^), the point at which network activity noticeably began degrading, is added to the individual connection weights between neurons. Although individual neurons in the network respond with different rates, the network is able to maintain persistent activity at many levels.

**Figure 5.**
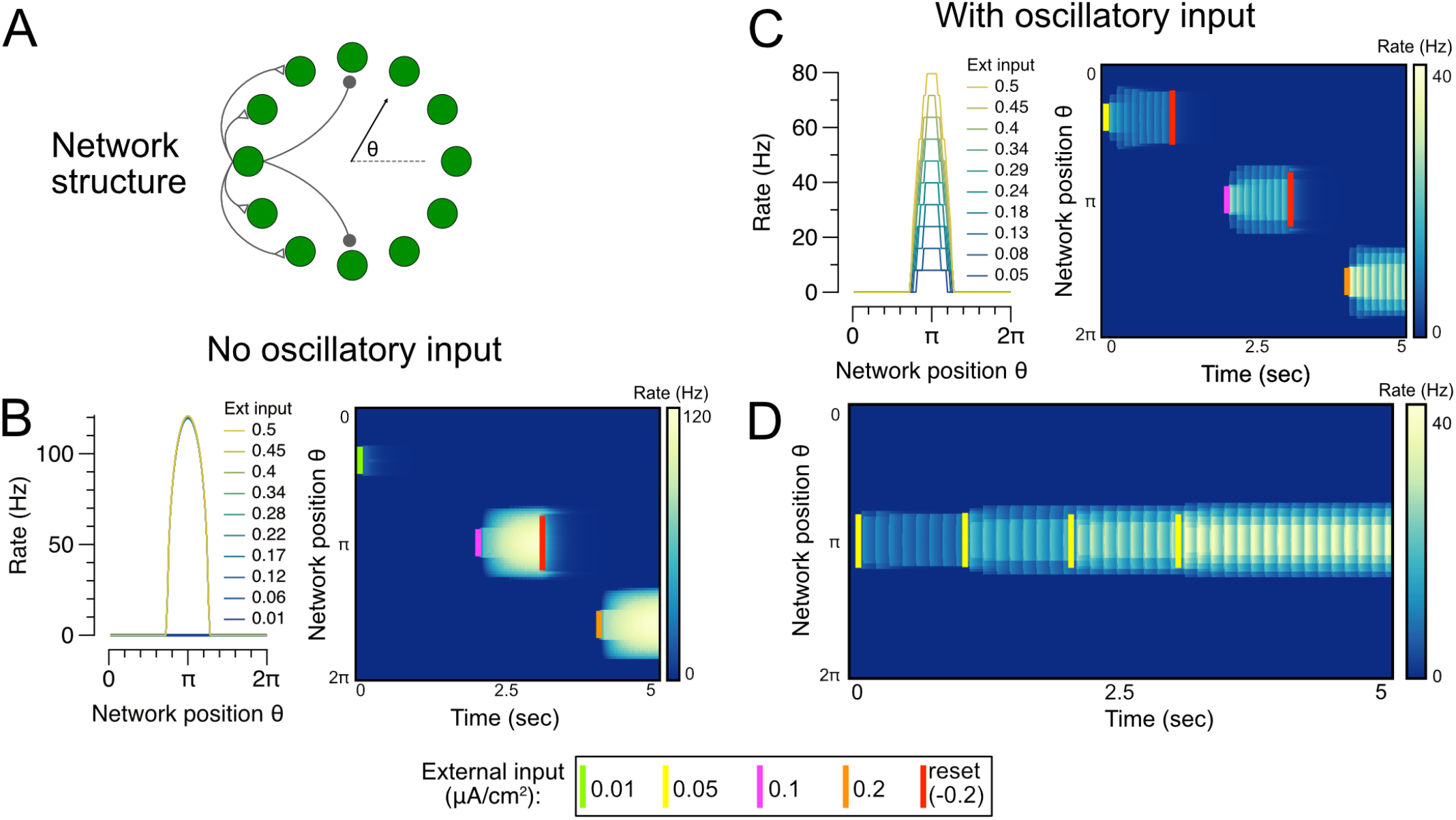
Maintenance of discretely graded bumps of persistent activity in a ring network. (A) Schematic of network structure. The spatial positions of neurons in the network are indexed by the angle theta from an arbitrary reference neuron. (B) Illustration of low-rate problem in a ring network of conductance-based spiking neurons without oscillatory input. Left, steady-state firing rate response of neurons in the network to input pulses of different amplitudes at locations centered around network position= *π*. Right, heatmaps illustrating the network’s temporal firing-rate responses to short pulses of inputs at network locations labeled by colored bars. The network is unable to maintain bump activity levels between the low and high fixed points. (C) Ring network with oscillatory input is able to maintain discretely graded bumps of persistent activity. (D) Temporal integration in the ring network. Short (100 ms) input pulses to the network are temporally integrated and stored in persistent activity.

**Figure 6.**
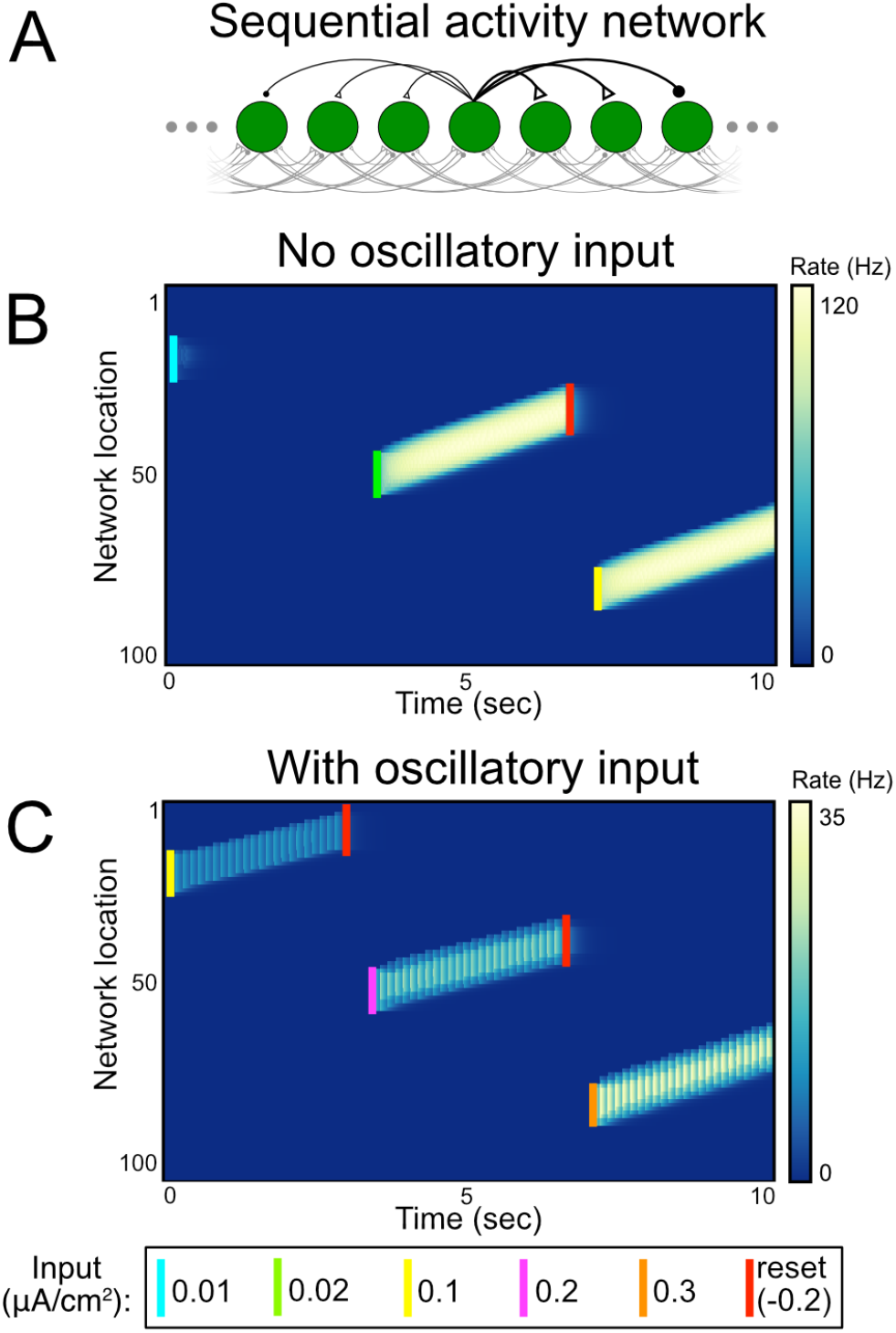
Generation of sequences of discretely graded activity in a network with asymmetric connectivity. (A) Schematic of network structure. Asymmetric connectivity underlies slow drift of activity bumps. (B) Illustration of low-rate problem in a sequential-activity network of conductance-based spiking neurons without oscillatory input. Drifting bumps of activity in the network initiated by short (100 ms) pulses (labeled by colored bars) exhibit only a single level of activity. (C) Sequential-activity network with oscillatory input is able to maintain drifting bumps with discretely graded levels of activity.

We first extended the demonstration of temporal integration, shown in Figure 3, to a spatially homogeneous (all-to-all) network composed of 1000 neurons (Fig. 4A,B). This permitted us to not only examine the systematic mistuning of weights shown in the autapse network, which produces identical results in the averaged activity of the 1000 neuron network (Methods), but also to examine the robustness to four different sources of noise and variability: input noise, in which each neuron in the network received independent exponentially filtered noise added to the subthreshold oscillatory drive (Fig. 4C); noise in the connection weights, in which each synapse in the network was initialized with added random noise (Fig. 4D); randomly shuffled phases of the subthreshold oscillatory drive, in which each neuron received an oscillatory signal whose phase was randomly picked from a uniform distribution on [0, 2**π**) at initialization (Fig. S2A); and noisy oscillation frequency and amplitude, in which the parameters of the subthreshold oscillatory drive underwent noisy drift (given by an Ornstein-Uhlenbeck process) during the simulations (Fig. S2B). In all of these cases, the network was able to accurately maintain multi-level persistent activity despite moderate perturbations. Figures 4C-D and Figure S2B illustrate the conditions for which the magnitude of the perturbations began to adversely affect network performance – for noise less than this amount, persistent activity was accurately maintained over a timescale of seconds, whereas larger noise levels led to progressively larger drifts of activity.

Next we demonstrate that a similar temporal integration of inputs can also occur in spatially structured networks. We consider a classic “ring model” architecture commonly used to model spatial working memory tasks in which stimuli can be presented at any of various locations arranged in a circular (ring-like) layout. The model consists of a ring of neurons with local excitatory connectivity and functionally wider inhibitory connectivity (Fig. 5A, Methods). Such models can generate persistent activity at any spatial location along the ring, but typically have only a binary “on-off” representation at a given spatial location (Fig. 5B). When we added an oscillatory input stimulus to such a ring model, the network could store multiple, discretely graded levels of activity at any spatial location (Fig. 5C) and could temporally integrate location-specific inputs into discretely graded levels (Fig. 5D). While the spatial memory (bump attractor) networks proposed in (38–40) are capable of generating graded persistent activity, the network presented here represents, to our knowledge, the first spatial memory network to encode multi-level activity without requiring an exact tuning condition.

Recent studies have shown that memory activity during a delay period also may take the form of a sequence of activity that spans the delay (41–43). Models of such activity typically generate chain-like patterns of activity that attain only a single, stereotyped level of firing rate. Consistent with this, when we constructed a network with a chain-like architecture (Fig. 6), we found that, in the absence of oscillatory input, the sequential network activity either quickly decayed when the initial stimulus amplitude was too small or converged to a single saturated level of activity for larger stimuli (Fig. 6B). By contrast, in the presence of a subthreshold oscillatory input, the network could exhibit sequential activity with discretely graded amplitudes for the same pattern of input (Fig. 6C). Thus, as in the persistently active networks, the oscillatory sequential memory network could encode multiple discretized stimulus levels.

In summary, this work demonstrates a simple mechanism by which oscillatory input to a memory network can transform it from storing only binary amplitudes to maintaining discretely graded amplitudes of persistent activity. Memory networks using this mechanism require a cellular, synaptic, or network process that can span the period of the oscillation, suggesting a possible tradeoff in memory storage: higher frequency oscillations do not require long timescale processes to span the oscillation cycle, but due to their short period may only store one or a few values (Figure 2F); lower frequency oscillations could store more items, but require a process with longer timescale to bridge the troughs occurring in each cycle. Our work complements traditional attractor models of working memory that typically fall into two classes: bistable models that robustly maintain two levels of activity (Figures 1B and 2B) and continuous attractor models that can maintain nearly analog storage of memory but require very precise tuning of connection weights (Figures 1D and 3C). Our model represents an intermediate possibility with relatively moderate tuning requirements (Fig. 3B) and a discretely graded set of response levels. Previous work (44, 45) has suggested how multiple, spatially distinct bistable processes in a cell can be coupled together to form multiple stable levels of firing activity; here we demonstrate a complementary mechanism for forming multi-stable representations that relies on temporal, rather than spatial, patterning of inputs. Altogether, this work suggests a potential mechanism by which oscillatory activity, which is commonly observed during working memory tasks, may expand short-term memory capacity.

## Methods

The Wang-Buzsaki model neuron used for most spiking neuron simulations in this paper is based upon the original model described in (34). Below, we show the equations for the dynamical variables most relevant to the maintenance of discretely graded persistent activity. The full model equations are included in the Supporting Information. The membrane potential of the Wang-Buzsaki neuron obeys the current balance equation:

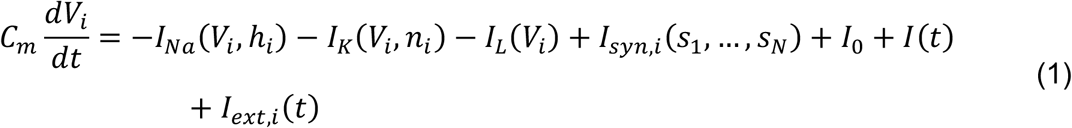

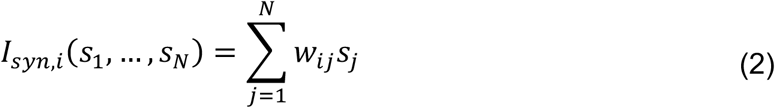

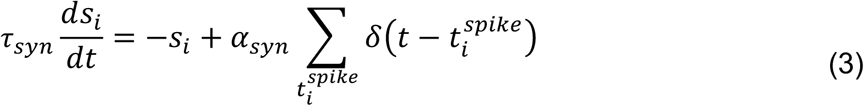

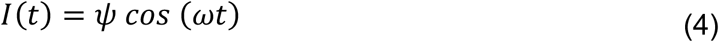

where *h* and *n* are time-varying channel variables (Supporting Information). The parameter values used are specified in Tables S1 and S2. The Wang-Buzsaki neuron receives several sources of inputs: (1) *I*_*syn,i*_(*s*_1_, …, *s*_*N*_) represents recurrent feedback to neuron *i*, the strength of which is determined by a weight matrix *w*_*ij*_ defining the strength of the connection from neuron *j* to neuron *i*, (2) *I*_0_ is a constant current that shifts the resting potential, and could represent tonic background input or intrinsic currents not explicitly modeled, (3) *I*(*t*) is the external oscillatory input (*I*(*t*) = 0 for models with no oscillatory input), and (4) *I*_*ext,i*_(*t*) represents the external inputs to be accumulated and stored by the memory network. To calculate spike times in equation 3, we used the time of the peak of the action potential, with only action potentials exceeding a voltage of 0 mV included. Integration was performed numerically using the fourth order Runge-Kutta method with a time step Δ*t* = 10^−2^ ms.

In the single neuron case, there is a single recurrent synaptic weight, *w* [*uA/cm*^2^]. Values for all simulation parameters are included in Table S2. In Figures 4-6, we study three different network architectures composed of Wang-Buzsaki neurons: an all-to-all connectivity (Figure 4), a ring structure (Figure 5), and a directed structure (Figure 6).

The all-to-all connected networks of Figure 4 are composed of 1000 Wang-Buzsaki neurons. Figures 4B,C implement a network with uniform connection strengths 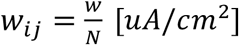. Figure 4D implements a network in which these uniform connection strengths have been perturbed by adding static Gaussian noise of mean zero independently to each connection. Exponentially filtered temporally white noise (Ornstein-Uhlenbeck process) input was implemented in the network illustrated in Figure 4C; for each neuron *i*, the additive noise was given independently by:

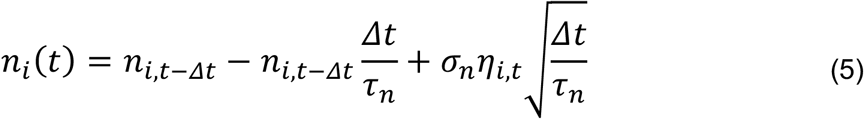

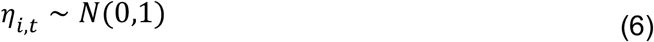

where *σ*_*n*_ is the standard deviation of the noise.

For the ring connectivity structure in Figure 5, the connection strength from neuron *j* to neuron *i* is described by:

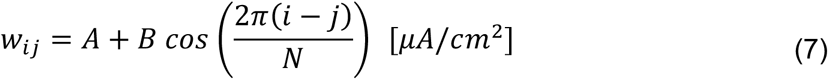

The directed structure illustrated in Figure 6 resembles the ring structure, but results in a drift of the ‘activity bump’ in one direction. The connection strength from neuron *j* to neuron *i* in this case is defined by:

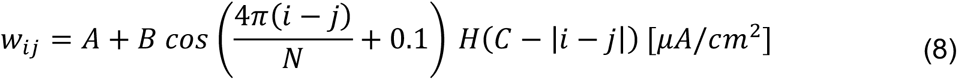

where *H* is the Heaviside (step) function and *C* controls the spatial extent of the connectivity.

### Comparison to linear spiking autapse model

In Figure 3, we compare the robustness of discretely graded persistent activity of the phase-locking nonlinear spiking model described above, to that of a spiking autapse model in which analog persistent activity is enabled by excitatory feedback that is tuned to offset the intrinsic decay of activity. The autapse model is described in detail in (37); equations describing the dynamics of the model are included in the Supporting Information.

### Simple rate model

The equation for the simple rate model implemented for Figure 1 is given in Figure 1A. The nonlinear term used for Figure 1B is:

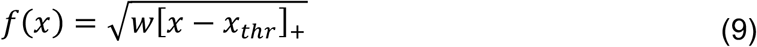

with *w* = 75, *x*_*thr*_ = 10 [*Hz*].

## Acknowledgements

We thank the 2016 Methods in Computational Neuroscience course (supported by NIH grant R25 MH062204 and the Simons Foundation) at the Marine Biological Laboratory, where this collaboration was started. Steve Luck provided helpful comments on the manuscript. This work was supported by NIH grants R01EY027036 (M. Goldman, B. Lankow) and U19NS104648 (M. Goldman, B. Lankow), NSF Graduate Research Fellowship DGE-1256082 (K. Champion), NSF grant DMS 1951099 (B. Ermentrout), and Swiss National Science Foundation no. 200020 184615 (O. Gozel).

## Supporting Information

### 1 Model neurons and networks

In this supplement, we provide a detailed description of the neuronal and network models used in this work. The multi-stable networks with oscillatory input make use of a conductance-based, single-compartment model neuron described in [3], which we refer to here as the Wang-Buzsaki (WB) neuron. The equations and parameter values governing the intrinsic response properties of this model, in which spiking is generated through Hodgkin-Huxley-type Na^+^ and K^+^ voltage-dependent ion currents, are described in section 1.1 below. In section 1.2, we describe the network models based on oscillatory input. In section 1.3, we describe the approximately linear autapse model, adapted from [1], whose robustness to detuning of the autapse strength was compared to that of the WB autapse model.

#### 1.1 Description of the Wang-Buzsaki neuron model

The membrane potential of the WB neuron obeys the current balance equation

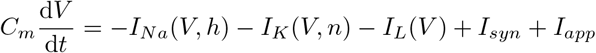

where the kinetics are as described in [3]:

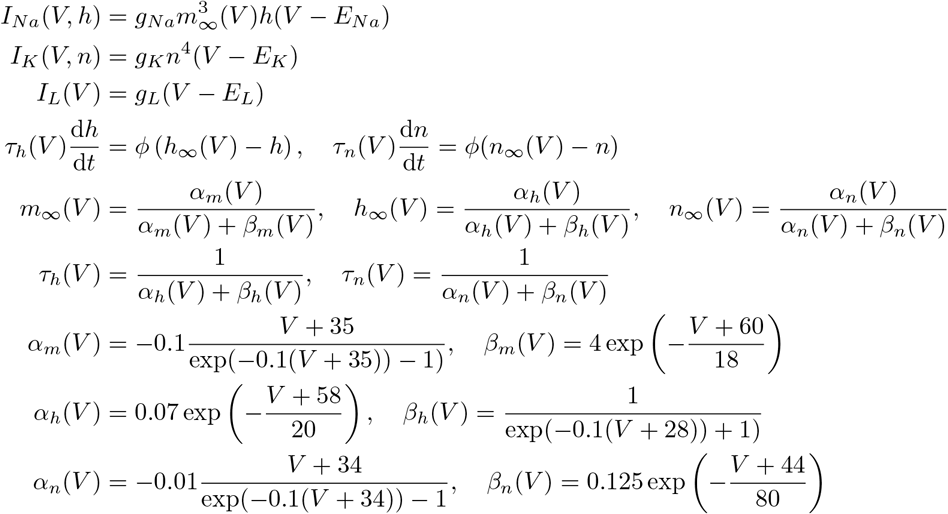

The synaptic and applied currents, *I*_*syn*_ and *I*_*app*_, respectively, are specified for different model architectures in the following sections. The common parameters used for all simulations involving WB model neurons are specified in Table 1. We defined the spike times as occurring when the membrane potential first decreased (d*V*/d*t* < 0) after crossing the spike threshold of 0 mV. Integration was performed numerically using the fourth-order Runge-Kutta method with time step Δ*t* = 10^−2^ ms.

**Table 1:**
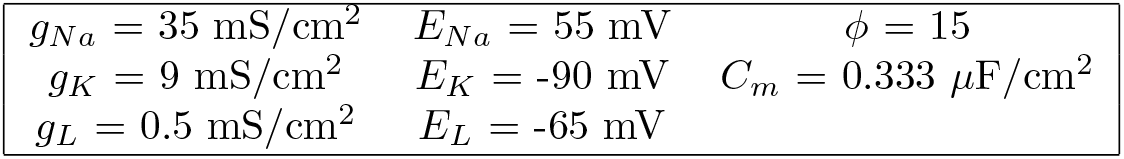
Common parameters of all WB neuron simulations

|  |  |  |
| --- | --- | --- |
| $g_{Na} = 35 \text{ mS/cm}^2$ | $E_{Na} = 55 \text{ mV}$ | $\phi = 15$ |
| $g_K = 9 \text{ mS/cm}^2$ | $E_K = -90 \text{ mV}$ | $C_m = 0.333 \text{ } \mu\text{F/cm}^2$ |
| $g_L = 0.5 \text{ mS/cm}^2$ | $E_L = -65 \text{ mV}$ | |

#### 1.2 Networks of Wang-Buzsaki model neurons

The different network architectures used in this work are described in the following subsections. The simplest of these, a single model neuron recurrently connected to itself (the autapse model), is described in section 1.2.1. In section 1.2.2, we describe the multi-neuron cases of all-to-all connected networks, a ring network, and a network that exhibits sequential drift of a bump of activity. The parameters specific to the autapse network simulations are listed in Table 2, and those specific to multi-neuron networks are listed in Table 3. Equations and parameter values for the stochastic simulations of multi-neuron networks are provided in Section 2.

**Table 2:** Parameters specific to the single-neuron (autapse) networks

| Non-oscillatory input | Oscillatory input |
| --- | --- |
| $w_{syn} = 1 \mu\text{A}/\text{cm}^2$ | $w_{syn} = 5.5 \mu\text{A}/\text{cm}^2$ |
| $I_0 = 4.005 \mu\text{A}/\text{cm}^2$ | $I_0 = 3.515 \mu\text{A}/\text{cm}^2$ |
| $\psi = 0 \mu\text{A}/\text{cm}^2$ | $\psi = -0.5 \mu\text{A}/\text{cm}^2$ |
| $\tau_{syn} = 150 \text{ ms}$ | $\tau_{syn} = 150 \text{ ms}$ |
| | $\omega = 0.05 \text{ ms}^{-1}$ (unless specified otherwise) |

**Table 3:** Parameters specific to the multi-neuron networks of WB neurons

| Non-oscillatory input | Oscillatory input |
| --- | --- |
| $w_{syn} = 1 \mu\text{A}/\text{cm}^2$ (unless specified otherwise)<br>$\tau_{syn} = 150 \text{ ms}$<br>$I_0 = 4.01 \mu\text{A}/\text{cm}^2$<br>$\psi = 0 \mu\text{A}/\text{cm}^2$ | $w_{syn} = 5.5 \mu\text{A}/\text{cm}^2$ (unless specified otherwise)<br>$\tau_{syn} = 150 \text{ ms}, 80 \text{ ms}$ (Figure S1C), $30 \text{ ms}$ (Figure S1A,B)<br>$I_0 = 3.515 \mu\text{A}/\text{cm}^2$<br>$\psi = -0.5 \mu\text{A}/\text{cm}^2$<br>$\omega = 0.05 \text{ ms}^{-1}$ ($\theta$ frequency, unless specified otherwise) |

##### 1.2.1 Simplest network example: the autapse

The autapse model consisted of a WB model neuron recurrently connected to itself and was described as follows:

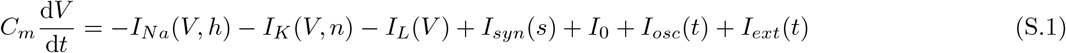

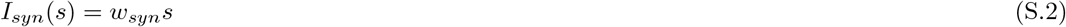

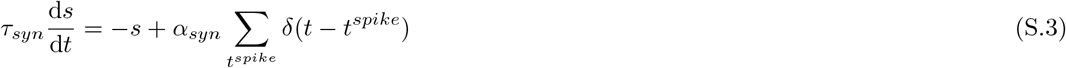

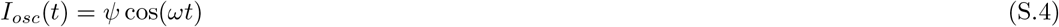

The neuron receives several sources of inputs: a recurrent synaptic feedback current *I*_*syn*_*(s)*, the strength of which is determined by *w*_*syn*_; a DC current offset *I*_0_; an oscillatory input current *I*_*osc*_*(t)* (set to zero for simulations with no oscillatory input); and a current corresponding to the external stimulus *I*_*ext*_*(t)*. The parameters specific to oscillatory and non-oscillatory simulations are listed in Table 2. *α*_*syn*_ *=* 1.0 ms for all simulations.

##### 1.2.2 Multi-neuron networks

For the networks composed of multiple WB neurons, each individual neuron *i* is governed by the same equations as for the single neuron case, except that the synaptic current input *I*_*syn*_,_*i*_*(s*_1_,…, *s*_*N*_) depends on the activity of all other neurons *j* of the network through the synaptic weights *w*_*ij*_:

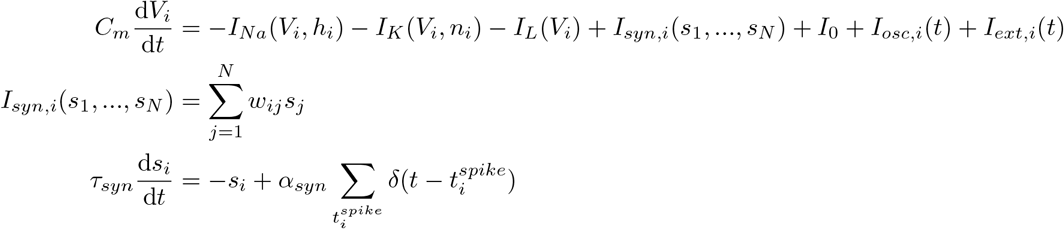

For all networks presented in this paper, we used a value of *α*_*syn*_ = 1.0 ms.

###### Fully connected network

In Figure 4 and Figures S1-S2, we implemented fully connected (all-to-all) networks of *N =* 1000 WB neurons. Except for the stochastic simulations described in Section 2, the connection strength from neuron *j* to neuron *i* was given by:

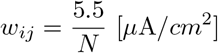

###### Ring network

For the ring network illustrated in Figure 5 (*N* = 100 neurons), the connection strength from neuron *j* to neuron *i* was described by a symmetric, rotationally invariant connection matrix:

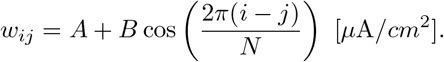

*A =* −0.024 [*μ*A/*cm*^2^] and *B* = 0.1515 [*μ*A/*cm*^2^] for the non-oscillatory case, and *A =* −0.54 [*μ*A/*cm*^2^] and *B* = 0.909 [*μ*A/*cm*^2^] for the oscillatory case.

###### Sequential activity network

In Figure 6, we illustrate an asymmetrically connected network (*N =* 100 neurons), which resembles the ring network, but whose asymmetry enables a well-controlled drift of the bump attractor in one direction. The connection strength from neuron *j* to neuron *i* was defined by:

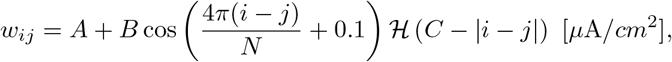

where ℋ denotes the Heaviside (step) function. *C =* 38 for all simulations. *A* = −0.18[*μ*A/*cm*^2^] and *B* = 0.303 [*μ*A/*cm*^2^] for the simulations with no oscillatory input, and *A* = −1.08 [*μ*A/*cm*^2^] and *B* = 1.818 [*μ*A/*cm*^2^] for the simulations with oscillatory input.

#### 1.3 Modified Seung autapse model

In Figure 3, we simulated a modified version of the model described in [1] (referred to here as the Seung autapse model) in order to compare its robustness to that of the WB autapse model with oscillatory input. The intrinsic neuron parameters are based on the model introduced by Shriki and colleagues [2], which exhibits an approximately linear f-I curve due to the inclusion of an A-type potassium current (*I*_*A*_(*V*_*k*_, *b*_*k*_) in the equations below). The model receives feedback current through a synapse onto itself (autapse). The model equations are:

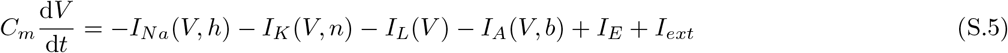

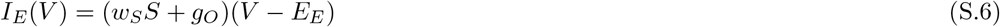

where the kinetics are as described in [1]:

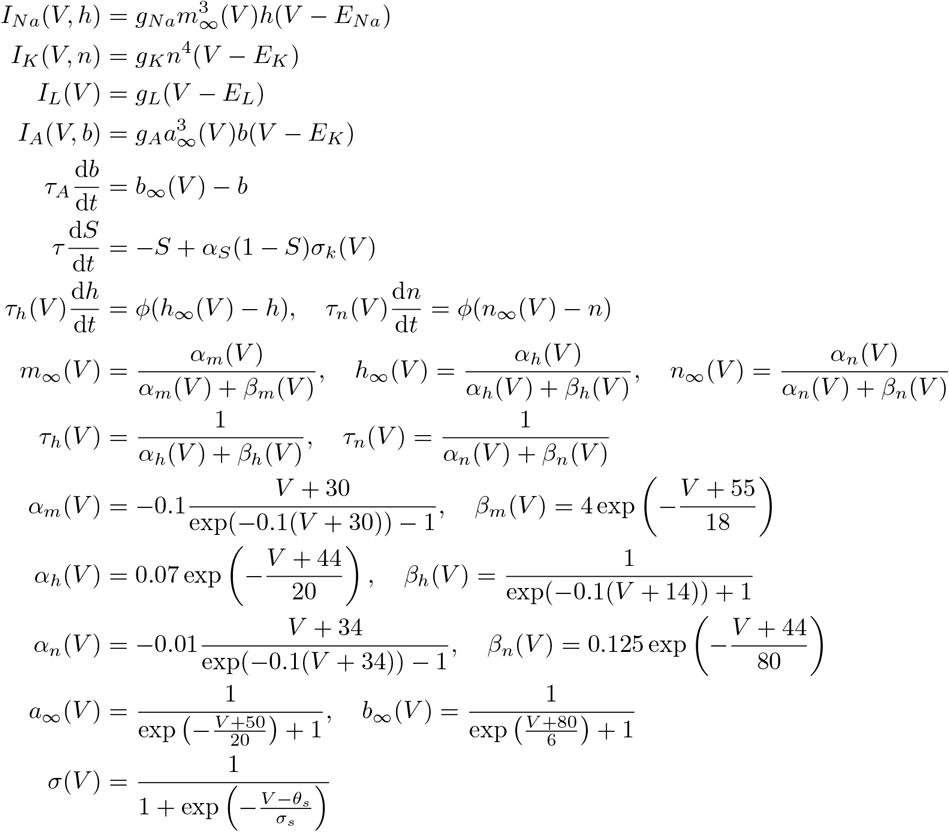

Individual parameters are listed in Table 4. Spike times are defined as the times of the downward zero crossings of the membrane potential after crossing a threshold of 20 mV. Integration was performed numerically using the fourth order Runge-Kutta method with time step Δ*t* = 2 · 10^−3^ ms.

**Table 4:**
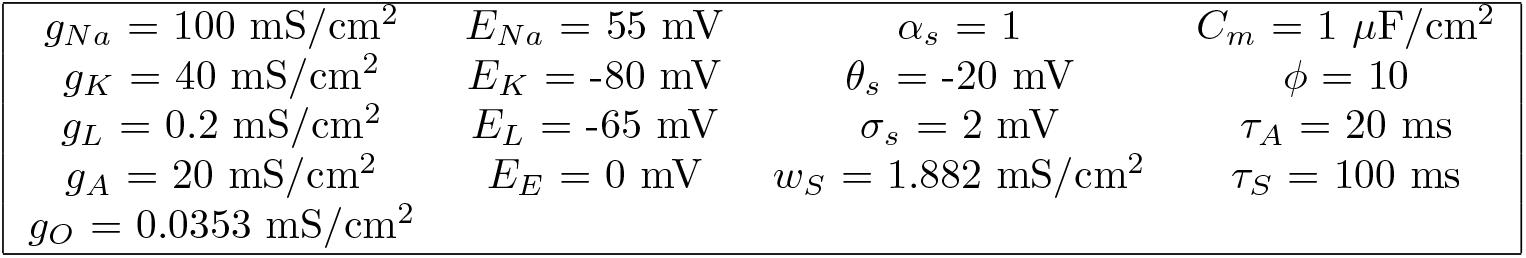
Parameters of the modified Seung autapse model

|  |  |  |  |
| --- | --- | --- | --- |
| $g_{Na} = 100 \text{ mS/cm}^2$ | $E_{Na} = 55 \text{ mV}$ | $\alpha_s = 1$ | $C_m = 1 \text{ } \mu\text{F/cm}^2$ |
| $g_K = 40 \text{ mS/cm}^2$ | $E_K = -80 \text{ mV}$ | $\theta_s = -20 \text{ mV}$ | $\phi = 10$ |
| $g_L = 0.2 \text{ mS/cm}^2$ | $E_L = -65 \text{ mV}$ | $\sigma_s = 2 \text{ mV}$ | $\tau_A = 20 \text{ ms}$ |
| $g_A = 20 \text{ mS/cm}^2$ | $E_E = 0 \text{ mV}$ | $w_S = 1.882 \text{ mS/cm}^2$ | $\tau_S = 100 \text{ ms}$ |
| $g_O = 0.0353 \text{ mS/cm}^2$ | | | |

### 2 Stochastic simulations

For the multi-neuron networks, we considered the effects of several forms of stochasticity: simulations with time-varying random noise in the inputs (section 2.1), simulations with “frozen” (not time-varying) noise in the synaptic weights of the network (section 2.2), and simulations in which the phases of the oscillatory external input to different neurons were randomized (section 2.3). We describe our methods for each of these cases below.

#### 2.1 Networks with time-varying noise in their inputs

In Figure 4C and Figure S2B, we simulated fully connected networks of 1000 WB neurons with two different types of time-varying noise: the addition of exponentially filtered Gaussian noise, *n*_*i*_*(t*), to the input (Figure 4C), or the inclusion of multiplicative filtered noise in the amplitude and frequency parameters (*ψ*(*t*) and *ω*(*t*), respectively) of the oscillatory input *I*_*i*_(*t*) (Figure S2B). In all cases, the noise was independent for each neuron in the network. The resulting models are described by:

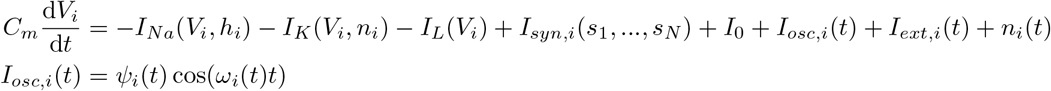

where *n*_*i*_*(t), ψ*_*i*_(*t*), and *ω*_*i*_(*t*) vary in time, as described in the sections below.

##### Noisy external input *n*_*i*_(*t*)

For the simulations illustrated in Figure 4C, exponentially filtered white noise (an Ornstein-Uhlenbeck process) input was implemented for each individual neuron *i* as:

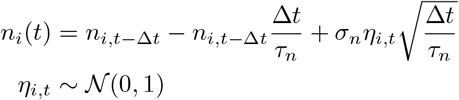

with *σ*_*n*_ *=* 0.05 *μ*A ms^0.5^/cm^2^.

##### Noisy oscillatory parameters *ψ*_*i*_(*t*)and *ω*_*i*_(*t*)

For the simulations illustrated in Figure S2B, the amplitude (*ψ*_*i*_) and frequency (*ψ*_*i*_) of the oscillatory input to each neuron evolved over time through multiplicative noise chosen independently for each neuron *i* of the network:

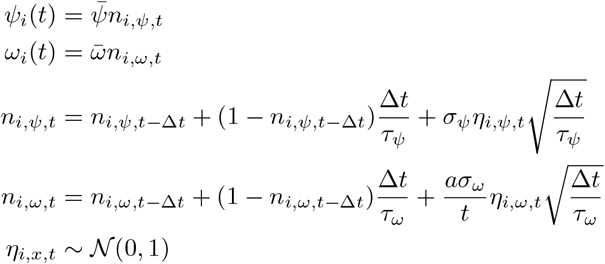

with parameters 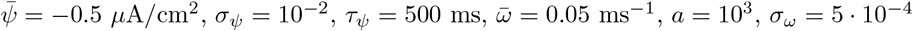, and *τ*_*ψ*_ = 500 ms.

#### 2.2 Networks with (frozen) noise in their synaptic weights

In the simulations illustrated in Figure 4D, the synaptic weights of the networks were initialized by independently adding random noise to the uniform synaptic weights. The random values were independently drawn from a normal distribution with *μ* = 0 [*μ*A/cm^2^] and standard deviation *σ* = 0.055 [*μ*A/cm^2^]. The value of the standard deviation was chosen to correspond to approximately where the network’s ability to maintain multi-level persistent activity became significantly degraded. This can be thought of as an approximate tolerance value of the 1000-neuron network to variation in the synaptic weights.

#### 2.3 Networks with randomized phases of oscillatory input

In Figures S1-S2, we demonstrate how the distribution of relative phases of oscillatory inputs affects the synaptic time constant needed to maintain multi-level persistent activity. The individual phases of the inputs to the network illustrated in Figures S1B and S2A were initialized by drawing randomly and independently from a uniform distribution on [0, 2*π*). In Figure S1C, the phases were initialized by drawing randomly and independently from a von Mises distribution with *μ* = 0 and concentration parameter *k* = 1.0. In all cases, the networks consisted of 1000 homogeneously connected WB neurons.

### 3 Parameters specific to simulations

Any parameter values used to generate figures included in the text or Supporting Information that were not stated above or in the figures are listed below.

**Figure 1**

All external input durations were 50 ms. The amplitudes of the pulses used to generate Figure 1C were [10 (green), 30 (purple), 40 (orange), 45 (red), 60 (blue)] Hz. The times of stimulus onset in Figure 1E were [100, 2000, 4000, 6000, 8000] ms. The amplitudes of the pulses used to generate Figure 1E (orange traces) were [70, 70, 70, -20, -20] Hz. The amplitudes of the pulses used to generate Figure 1E (blue traces) were [50, 50, 50, -40, -40] Hz. The amplitudes of the pulses used to generate Figure 1E (red traces) were [30, 30, 30, -130, -130] Hz.

**Figure 2**

All external input durations were 100 ms and the time of stimulus onset was t=100 ms. The amplitudes of the pulses used to generate Figure 2B were [0.011 (green), 0.015 (purple), 0.0175 (orange), 0.13625 (red), 0.2 (blue)] *μ*A/cm^2^. The amplitudes of the pulses used to generate Figures 2D,E,F were [0.05 (green), 0.2 (purple), 0.3 (orange), 0.45 (red), 0.6 (blue)] *μ*A/cm^2^.

**Figure 3**

All external input durations were 100 ms for Figure 3A and 50 ms for Figure 3C; the times of stimulus onset were [100, 1100, 2100, 3100, 4100, 5100, 6100, 7100, 8100, 9100] ms. The amplitudes of the pulses used to generate Figure 3A were [0.2, 0.1125, 0.15, -0.225, 0.225, -0.225, -0.1125, 0.15, -0.15, 0.1125] *μ*A/cm^2^. The amplitudes of the pulses used to generate Figure 3C were [2.0, 0.75, 1.0, -1.5, 1.5, -1.5, -0.75, 1.0, -1.0, 0.75] *μ*A/cm^2^.

**Figure 4**

All external input durations were 100 ms and the times of stimulus onset were [100, 2000, 4000, 6000, 8000, 10000, 12000, 14000, 16000, 18000] ms. The amplitudes of the pulses used to generate Figure 4B were [0.1125, 0.125, 0.1375, 0.175, 0.175, 0.2, -0.175, -0.175] *μ*A/cm^2^. The amplitudes of the pulses used to generate Figure 4C were [0.1, 0.125, 0.125, 0.1375, 0.175, 0.175, 0.2, -0.175, -0.175, -0.175] *μ*A/cm^2^. The amplitudes of the pulses used to generate Figure 4D were [0.1, 0.125, 0.125, 0.125, 0.125, 0.125, 0.125, -0.175, -0.175, -0.175] *μ*A/cm^2^.

**Figure S1**

All external input durations were 50 ms and the times of stimulus onset were [125, 2125, 4125, 6125, 8125, 10125] ms. The amplitudes of the pulses used to generate Figure S1A were [0.05, 0.1, 0.15, 0.2, 0.25, 0.3] *μ*A/cm^2^. The amplitudes of all pulses used to generate Figure S1B were 0.03125 *μ*A/cm^2^. The amplitudes of all pulses used to generate Figure S1B were 0.075 *μ*A/cm^2^.

## 5 Supplementary figures

**Figure S1.**
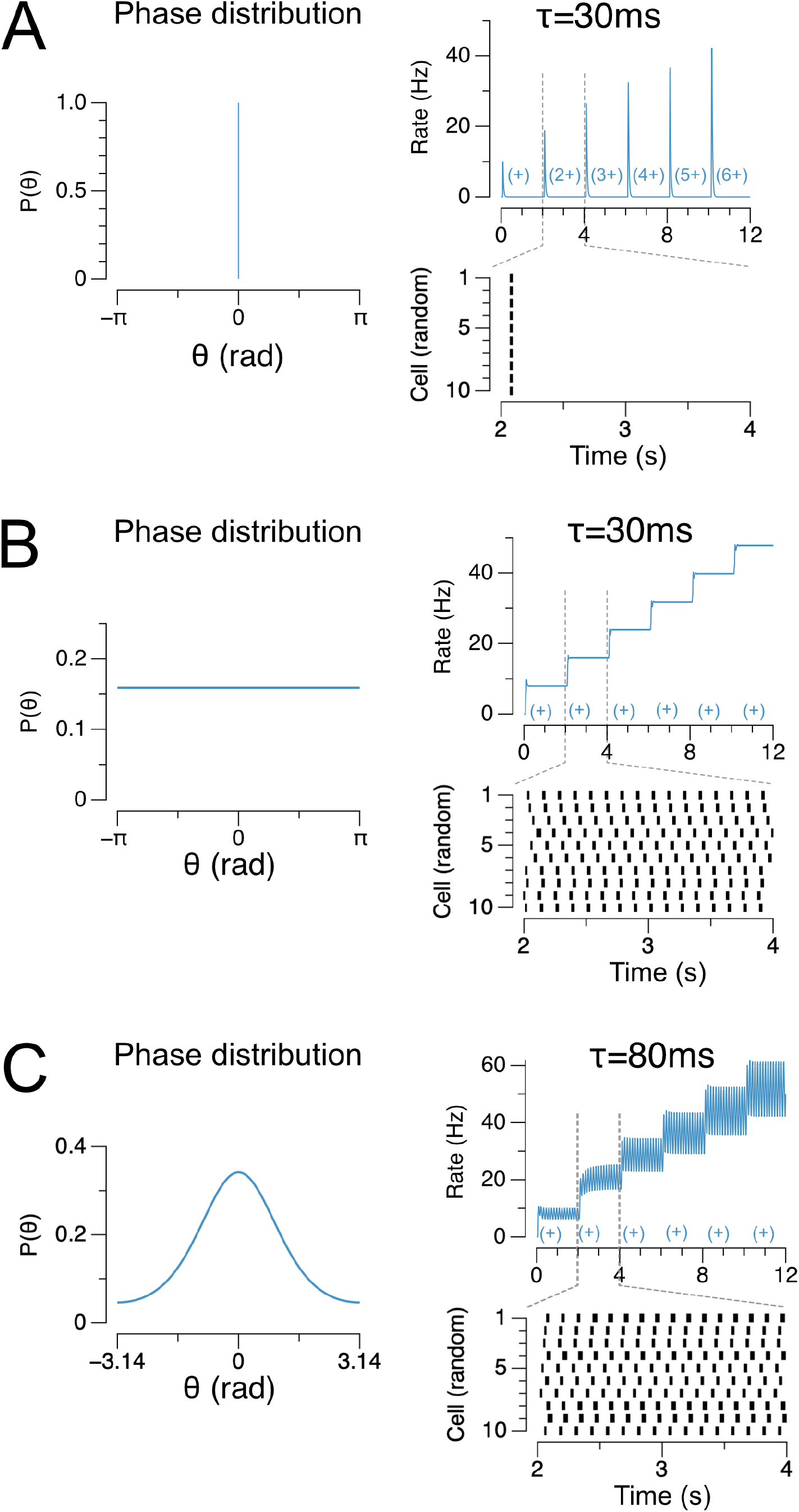
Heterogeneity in the phases of oscillations in the network enables discretely graded persistent activity using synaptic time constants that are shorter than the oscillation cycle. (A) Failure of network with perfectly aligned phases and shorter synaptic time constants to maintain persistent activity. When all oscillatory phases are perfectly aligned (left), the currents provided by the shorter time constant synapses (*τ* = 30 ms) cannot bridge the troughs of the oscillation cycles, and the network fails to maintain persistent firing (right). Times and sizes of external pulses of increasing magnitude are depicted along the x-axis with (n+) denoting a pulse of n times the magnitude of (+). (B) Distributed phases of oscillatory inputs enables the network to store discretely graded activity even with 30 ms synaptic time constants. When phases of the oscillatory inputs to the network are independently drawn from a uniform distribution (left), the relative spread of spiking responses in the network can bridge the oscillation cycles and enable discretely graded persistent activity. (C) Network response to a distribution of oscillatory inputs intermediate between the phase distributions illustrated in (A,B). Left, distributions were generated by sampling the phases from a Von Mises distribution (concentration parameter *k* = 1.0). Right, the network can maintain discretely graded persistent activity with synaptic time constants (*τ* = 80 ms shown here) typically used in studies of persistent activity.

**Figure S2.**
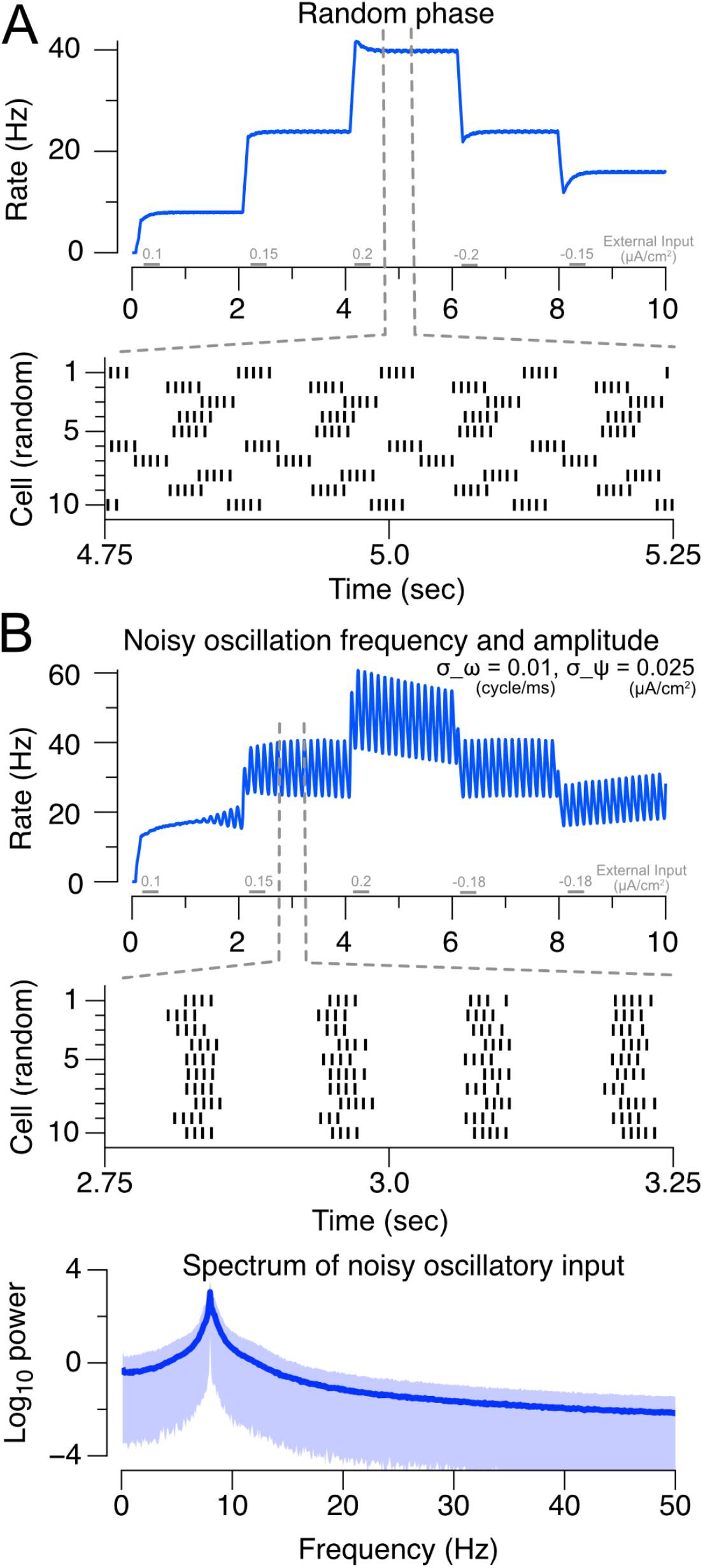
Robustness of persistent activity in a fully connected network of 1000 WB neurons receiving noisy oscillatory input or oscillatory input with randomized phase across neurons. (A) Mean firing rate across the population (top) and rasters of individual neuron responses (bottom) for a network in which the phase of oscillatory input is randomly chosen from a uniform distribution for each neuron. Input to the network is a sequence of brief (100 ms) input pulses (gray bars). (B) Mean firing rate across the population (top) and rasters of individual neuron responses (middle) to a sequence of brief input pulses in a network in which the values of the oscillatory parameters omega and psi vary randomly in time (Supporting Information). Bottom, power spectrum of the oscillatory input. Noise is independent (uncorrelated) across neurons of the network.

